# Investigating the Neural Correlates of Processing Basic Emotions: A Functional Near-Infrared Spectroscopy (fNIRS) Study

**DOI:** 10.1101/2023.08.08.551979

**Authors:** Gülnaz Yükselen, Ozan Cem Öztürk, Gümüş Deniz Canlı, Sinem Burcu Erdoğan

## Abstract

Emotion regulation, a fundamental aspect of human functioning, involves the ability to monitor, evaluate, and modify emotional responses. Understanding the neural mechanisms underlying emotion regulation holds significant implications across various disciplines, including psychology, neuroscience, and clinical psychiatry. This study aims to explore the neural correlates of emotion regulation using functional near-infrared spectroscopy (fNIRS) with a specific focus on the prefrontal cortex (PFC). fNIRS, a non-invasive and portable brain imaging technology, offers an excellent opportunity to investigate real-life emotion processing with high temporal resolution. Twenty participants underwent an experimental protocol where they viewed emotional pictures from the International Affective Picture System (IAPS) database, varying in valence (positive and negative) and arousal (high and low). fNIRS data were collected during the picture presentation, and the hemodynamic responses in the PFC were analyzed. The findings demonstrated distinct spatiotemporal patterns of activation associated with different emotional states. Positive valence stimuli elicited higher hemodynamic activation in bilateral dorsolateral prefrontal cortex (DLPFC) and orbitofrontal cortex (OFC) regions when compared to negative valence stimuli. On the other hand, negative valence stimuli induced higher activation in the medial prefrontal cortex (mPFC) when compared to positive valence stimuli. Moreover, high arousal positive valence stimuli evoked higher activation in the left DLPFC region when compared to high arousal negative valence stimuli. These results shed light on the differential neural processing of positive and negative emotions within the PFC, supporting the notion of lateralized emotional processing. The study validates the feasibility of fNIRS for objectively capturing emotion-related neural activity, providing valuable insights for future applications in emotion recognition and affective brain-computer interfaces (BCIs). Understanding the neural basis of emotion regulation has significant implications for designing targeted interventions for individuals experiencing emotion dysregulation disorders. Additionally, the integration of fNIRS technology into affective BCIs may offer new possibilities for real-time emotion detection and communication in populations with communication challenges.

## 1. INTRODUCTION

In the past few decades, there has been a surge of interest in the unraveling of intricate mechanisms involved in the processing and regulation of fundamental emotions in the healthy adult brain (Strait & Scheutz, 2014). Emotion regulation encompasses the ability to monitor, evaluate, and modify our emotional responses and plays a fundamental role in human functioning (Morawetz et al., 2017). Understanding the underlying neural mechanisms of emotion regulation carries significant implications across various disciplines, including psychology, neuroscience, and clinical psychiatry. Furthermore, investigating the spatiotemporal patterns of neural correlates associated with emotion processing and regulation bears profound consequences for the advancement of state-of-the-art affective Brain-Computer Interfaces (BCIs) (Hoshi et al., 2011; Lorenzetti et al., 2018). These interfaces aim to decode cognitive and affective neural responses in real-time, thereby presenting new possibilities for objectively discerning emotions and preferences in individuals encountering challenges in verbal communication, such as those with dementia or locked-in syndrome (Fenton & Alpert, 2008; Sun et al., 2022).

To precisely assess the neural processing of emotions within real-life scenarios, an increasing demand exists for wearable and non-invasive sensors capable of providing reliable and real-time data. Among the various modalities of functional brain imaging currently available, functional near-infrared spectroscopy (fNIRS) has emerged as a promising instrument for this purpose. fNIRS encompasses an optical brain imaging technology that non-invasively captures alterations in the levels of oxygenated and deoxygenated hemoglobin, reflective of neuronal activity (Ferrari & Quaresima, 2012). In comparison to alternative methodologies such as electroencephalography (EEG) and functional magnetic resonance imaging (fMRI), fNIRS exhibits less susceptibility to motion artifacts and offers higher temporal resolution. Furthermore, it facilitates data acquisition within ecologically valid settings, making it well-suited for investigating the intricate landscape of emotion processing (Ferrari & Quaresima, 2012; Pinti et al., 2020; Quaresima & Ferrari, 2019; Tak & Ye, 2014). The portability, user-friendliness, and minimal interference associated with fNIRS systems further enhance their appeal for BCI designs.

The prefrontal cortex (PFC) stands as a pivotal brain region implicated in the regulation of emotions. The investigation of the underlying neural networks of emotion has consistently been a prominent area of research within neuroscience. The widely accepted ‘limbic system’ model, initially proposed by Maclean in 1949, postulates the involvement of various subcortical brain regions, including the ‘reptilian brain’, in the mechanisms governing emotions. This model suggests that emotions, from intricate emotional states to more primal ones like fear, originate from the limbic system. However, the PFC appears to be responsible for the processing and regulation of emotions (Balconi & Molteni, 2016; Doi et al., 2013). The PFC plays a pivotal role in the evaluation of emotional stimuli, the generation of appropriate emotional responses, and the modulation of emotional intensity. Extensive investigation utilizing various neuroimaging techniques, including fNIRS, consistently highlights the involvement of the PFC in the regulation of emotions (Glotzbach et al., 2011; Ozawa et al., 2019; Tak & Ye, 2014). By integrating fNIRS into affective BCIs, we stand to gain valuable insights into the specific contributions of the PFC in these intricate processes. The integration of fNIRS technology into affective BCI systems holds immense potential for shedding light on the specific contributions of the PFC in these intricate processes.

Nevertheless, before achieving such integration, it is crucial to determine whether fNIRS can capture distinct spatiotemporal patterns of activation in the PFC during the processing of different basic emotions. Valence (i.e., the extent to which an emotion is positive or negative) and arousal (i.e., the strength of parasympathetic response) represent fundamental dimensions of emotion that significantly shape our emotional experiences. Within this context, the International Affective Picture System (IAPS) database provides standardized pictures with predefined valence and arousal scores and offers a robust framework for designing experimental protocols that target mapping the neural correlates of processing emotional stimuli in terms of valence and arousal dimensions (Lang et al., 2008). Numerous studies investigated the hemodynamic correlates of emotion processing by fNIRS recordings while the subjects were exposed to pictures from the IAPS database including those conducted by Balconi & Molteni, (2016); Glotzbach et al., (2011); Herrmann et al., (2003); Ozawa et al., (2014). Despite the extensive research conducted on the PFC and its involvement in emotion processing, the specific functions of the PFC in this context remain uncertain. Balconi & Mazza (2010), proposed lateralization effects in PFC activation during emotional processing, with positive valence being more strongly associated with the left hemisphere and negative valence with the right hemisphere. Similarly, Balconi & Molteni (2016) found higher z-scores of cortical activation during the processing of negative stimuli in both hemispheres when compared to positive stimuli, with a stronger right hemisphere bias for negative pictures. However, other studies suggest a left hemisphere bias for negative emotions such as fear and anger compared to neutral conditions (Glotzbach et al., 2011; Herrmann et al., 2008). Moreover, PFC activation may not be solely determined by valence but could represent a dichotomy between approach and avoidance attitudes toward emotions (Davidson, 1995; Harmon-Jones, 2003).

Inconsistent findings also emerge from studies on PFC involvement in emotion regulation using functional neuroimaging. Some studies indicate a crucial role of the PFC in regulating emotions (Wager et al., 2008), while others propose that the ventromedial prefrontal cortex does not mediate the regulatory effect of physical contact on neural threat responses (Lee et al., 2012). Furthermore, increased activation in the dorsolateral and ventrolateral PFC, regions associated with emotion regulation, has been observed in individuals with post-traumatic stress disorder (Nicholson et al., 2017). These inconsistent results suggest the involvement of different PFC regions in distinct aspects of emotion regulation, necessitating further research to comprehensively comprehend the neural mechanisms underlying emotion regulation in diverse contexts (Morawetz et al., 2016).

Building upon these considerations, our study aims to explore the neural correlates of emotion regulation utilizing fNIRS, with a specific focus on the PFC. By investigating how different levels of valence (i.e., positive and negative) and arousal (i.e., high and low) influence hemodynamic responses in the PFC, we aim to identify unique spatiotemporal patterns associated with each dimension of emotion. The findings of this study will significantly contribute to our understanding of the underlying neural mechanisms of emotion regulation, particularly concerning valence and arousal. Furthermore, the insights gained from this research may have important implications for clinical applications, including the development of targeted interventions for individuals experiencing emotion dysregulation disorders.

## 2. MATERIALS AND METHODS

### 2.1. Participants

Twenty-three volunteer subjects with no recent history of neurological or psychiatric disorders participated in this study. Data from three participants were excluded from the analysis due to poor signal quality, leaving a final sample size of twenty participants (mean age: 22±2, 11 females) for data analysis. Prior to the experiment, all participants provided written informed consent in accordance with the latest Declaration of Helsinki. The study was approved by the local ethics committee of Istanbul Medipol University, Istanbul, Turkey.

### 2.2. Materials

The present study employed pictures from the IAPS database, a well-established tool widely utilized in affective neuroscience studies that target elucidation of the neurobiological mechanisms that underlie emotional responses. The selection of pictures was conducted based on specific criteria, involving the assessment of picture-specific valence and arousal scores. For the inclusion of pleasant pictures, a prerequisite was that the valence score of each picture fell within the top 20%, while for unpleasant pictures, the valence score fell within the bottom 20%. Additionally, the high arousal condition necessitated an arousal score within the highest 30%, whereas the low arousal condition required an arousal score within the lowest 30%. Consequently, the study design involved a total of 40 unpleasant pictures (comprising of 20 high arousal and 20 low arousal), 40 pleasant pictures (consisting of 20 high arousal and 20 low arousal), as well as 20 neutral pictures, adhering to the aforementioned selection criteria. Selected pictures were the following: Pleasant & High Arousal; 2216, 2347, 4220, 4599, 4626, 4626, 4668, 5623, 5629, 5833, 7330, 7502, 8030, 8163, 8180, 8185, 8370, 8380, 8461, 8496. Pleasant & Low Arousal; 1441, 1600, 1601, 1610, 1620, 1920, 2035, 2170, 2314, 2341, 2360, 2387, 2510, 2530, 5001, 5201, 5831, 5982, 7325, 7470. Unpleasant & High Arousal; 2370, 3053, 6021, 6313, 6520, 6831, 7380, 9040, 9075, 9163, 9183, 9252, 9300, 9410, 9413, 9423, 9635.1, 9910, 9921. Unpleasant & Low Arousal; 2205, 2301, 2456, 2750, 3300, 9041, 9046, 9180, 9220, 9265, 9280, 9291, 9330, 9342, 9395, 9470, 9471, 9561, 9830. Neutral; 7000, 7001, 7002, 7003, 7004, 7006, 7009, 7010, 7012, 7014, 7017, 7019, 7020, 7025, 7034, 7050, 7052, 7186, 7500, 7950.

**Table 1.**
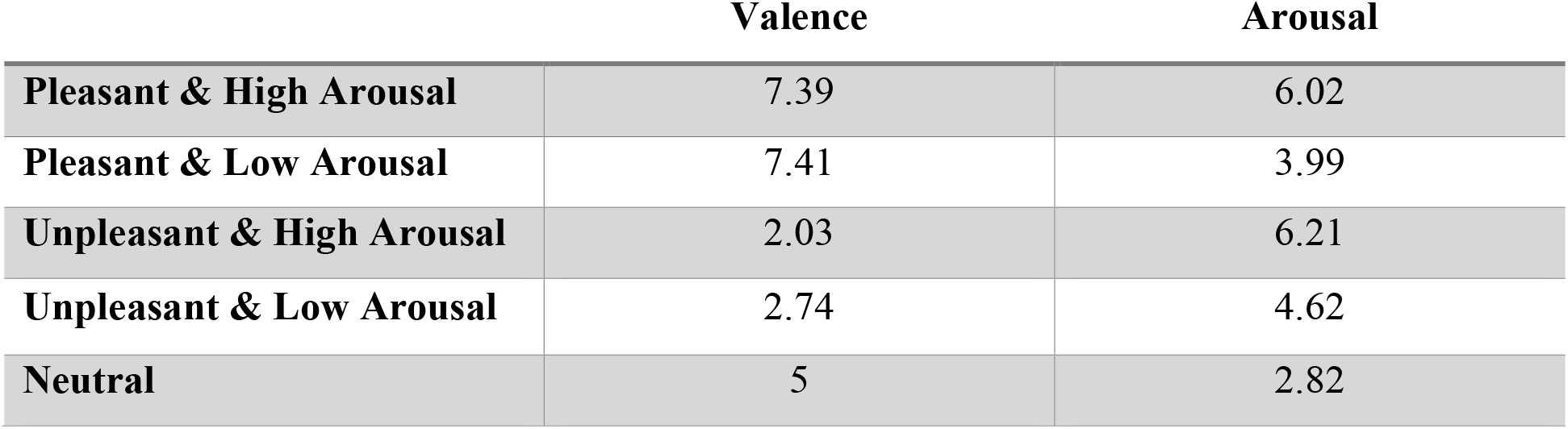
Valence and Arousal Ratings for Different Stimulus Conditions.

|  | Valence | Arousal |
| --- | --- | --- |
| Pleasant & High Arousal | 7.39 | 6.02 |
| Pleasant & Low Arousal | 7.41 | 3.99 |
| Unpleasant & High Arousal | 2.03 | 6.21 |
| Unpleasant & Low Arousal | 2.74 | 4.62 |
| Neutral | 5 | 2.82 |

### 2.3. Experimental Protocol

Participants were seated comfortably in a chair, positioned approximately 1m away from a computer screen in a quiet and dimly lit room. They were instructed to relax and minimize any unnecessary movements throughout the experiment. Prior to commencing the study, detailed information about the experiment was provided to each participant, and they were shown sample pictures for each experimental condition prior to the onset of the experiment. Participants were explicitly instructed to remain engaged in the task to the best of their ability and to focus their attention on the presented pictures.

To establish a baseline measurement, a 30-second period was allocated before the onset of presenting the stimuli blocks. Following this baseline measurement, the participants were presented with a series of pictures. The experiment consisted of 4 cycles of 5 consecutive blocks of each condition (i.e., high positive, low negative, low positive, high negative, and neutral) interleaved by 20-second rest periods. Each stimulus block consisted of 5 pictures of similar valence and arousal scores, each of which was presented on the screen for 5 seconds interleaved with 10 seconds of rest (Figure 1).

**Figure 1.**
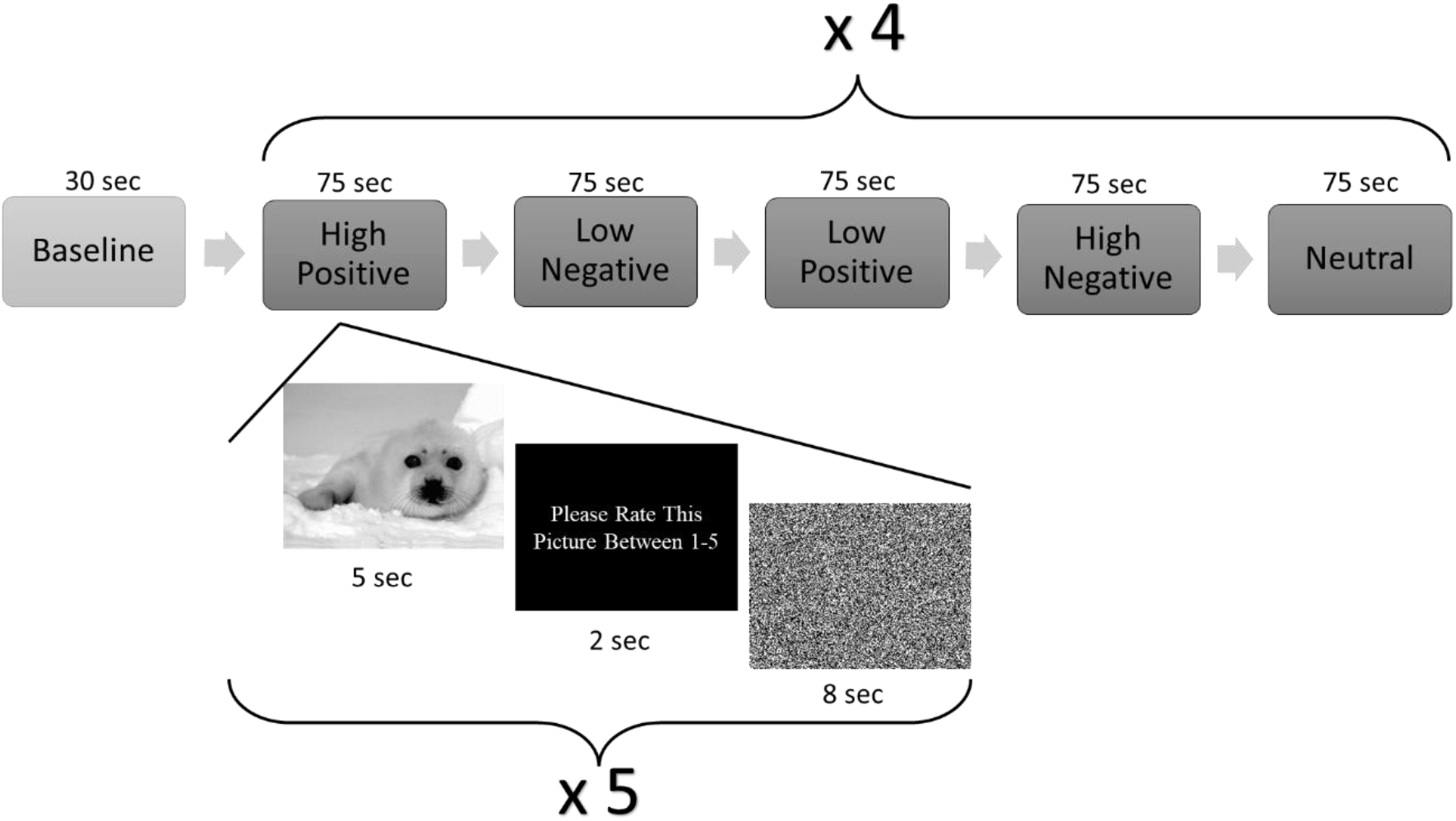
Experimental protocol. Participants underwent a 30-second baseline measurement before the presentation of stimuli blocks. The experiment comprised four cycles, each consisting of five consecutive blocks for each condition (High Positive, Low Negative, Low Positive, High Negative, and Neutral). Rest periods of 20 seconds were interleaved between the cycles. Each stimulus block included five pictures with similar valence and arousal scores, presented for 5 seconds each, interleaved with 10-second rest intervals.

### 2.4. fNIRS Data Acquisition

The hemodynamic data were collected using a NIRSport system (NIRx Medical Technologies, LLC, Berlin, Germany). The system utilized a specific configuration comprising of 22 channels, including 8 light sources emitting near-infrared light at wavelengths of 760 nm and 850 nm, and 7 detectors positioned over the forehead. The sampling rate of the signal was set at 7.81 Hz, and the channel forming sources and detectors were placed 3 cm apart.

In order to capture hemodynamic activity within the PFC region, the head probe was carefully positioned on the forehead, with the lower row of detectors placed slightly above the eyebrows. The arrangement of optodes and the distribution of channels can be visualized in Figure 2. Prior research has demonstrated that each light source within this probe configuration emits near-infrared light that has the ability to penetrate the scalp and reach the top 2-3mm of cortical gray matter tissue. This characteristic enables the investigation of various areas within the PFC, including the dorsolateral, orbitofrontal, and medial regions (Chance et al., 1998).

**Figure 2.**
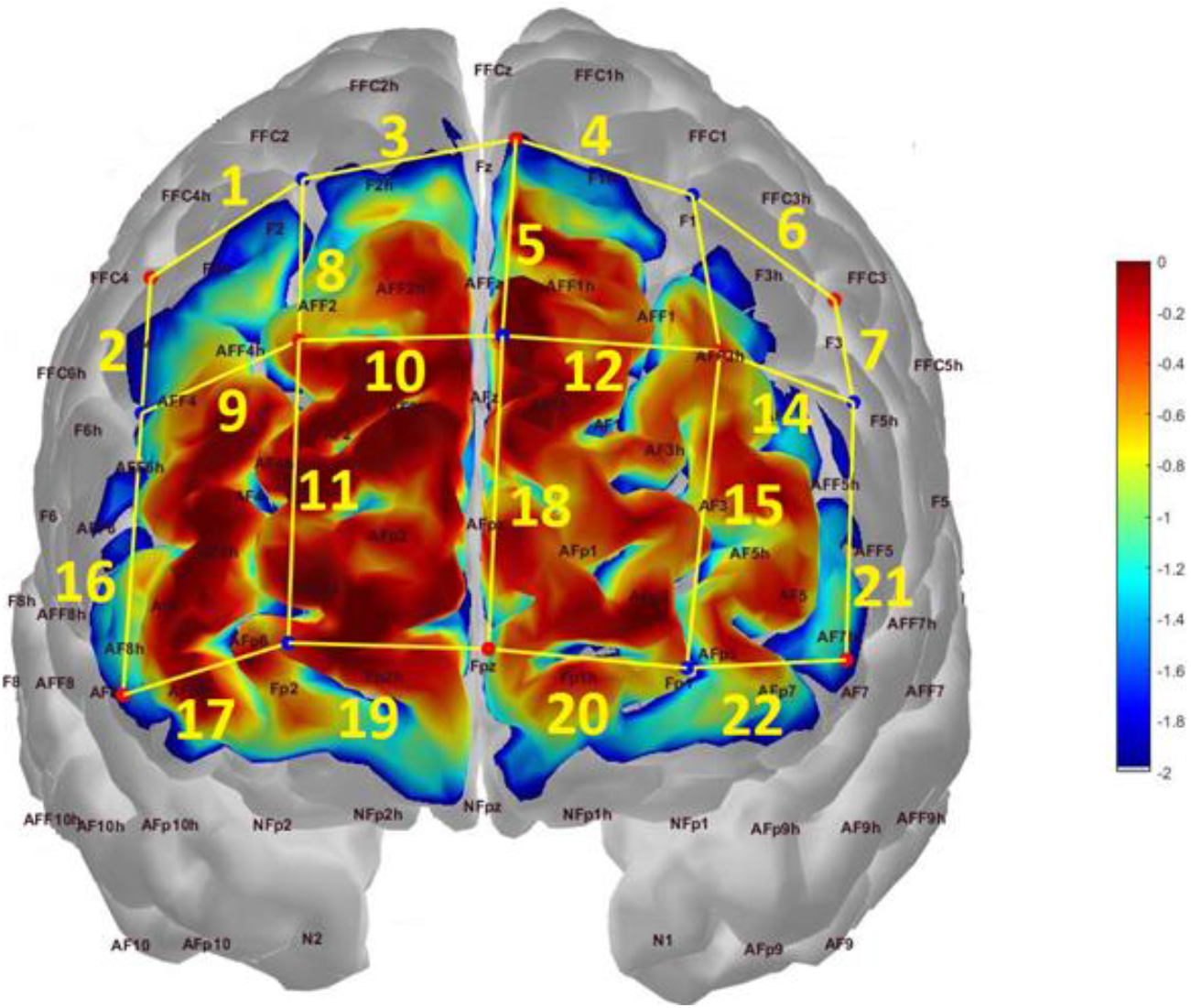
Probe configuration and photon sensitivity profile. Red dots indicate light sources, while blue dots represent detectors. Yellow lines connect adjacent source-detector pairs, forming channels labeled accordingly. The probe’s sensitivity to detecting brain hemodynamics is visualized as a heat map in log10 units. The color-coded heat map illustrates the varying levels of sensitivity across different regions of the brain.

Concentration changes in oxygenated hemoglobin (HbO) and deoxygenated hemoglobin (HbR) signals were determined utilizing the modified Beer-Lambert law with the two available wavelengths. This methodological approach facilitates the estimation of hemodynamic fluctuations occurring in the PFC region during the experiment.

### 2.5. Data Preprocessing and Feature Extraction

The preprocessing of the fNIRS signals was conducted using a combination of MATLAB scripts (Mathworks, Natick, MA, USA) and functions obtained from the Homer3 software (Huppert et al., 2009). The initial step involved visually examining the raw light intensity data. Subsequently, channels with poor signal quality were identified using the coefficient of variation (CV) method, as described in previous studies Da Silva Ferreira Barreto et al., (2020); Piper et al., (2014); Pollonini et al., (2016); Zimeo Morais et al., (2017). The CV was computed as a percentage using the formula CV (%) = 100 × (standard deviation of data)/(mean of data), where ‘data’ refers to the raw optical signals at 760 nm or 850 nm specific to each channel. Channels with raw light intensity signals exhibiting a CV above 7.5% were deemed heavily contaminated with unphysiological noise and were excluded from further analysis, following previous research (Hocke et al., 2018; Mutlu et al., 2020; Zimeo Morais et al., 2017). Additionally, subjects whose fNIRS data contained more than 5 channels of low signal quality were excluded from subsequent analysis, resulting in the exclusion of 3 subjects based on this criterion.

For the remaining subjects, motion artifacts in the data segments were identified for each channel using the hmrR_MotionArtifactByChannel function, utilizing specific parameters including tMotion, tMask, STDEVthresh, and AMPthresh, as described by Huppert et al., (2009). The raw intensity data were then converted into optical density (OD) data using the hmrR_Intensity2OD function. Any spikes and drifts observed in the channel OD data were initially corrected using the hmrR_MotionCorrectSplineSG function, with parameters p set to 0.99 and FrameSize_sec set to 10. The motion corrected OD data was further processed by converting them to HbO and HbR data using the hmrR_OD2Conc function, with the differential pathlength factor (DPF) set to 1 for each wavelength. Subsequently, the HbO and HbR data were subjected to bandpass filtering using a Butterworth filter embedded in the hmrR_BandpassFilt function, with cutoff frequencies set at 0.01-0.5 Hz. The remaining traces of motion artifacts were addressed by employing the correlation-based signal improvement method (CBSI) through the hmrR_MotionCorrectCbsi function. After bandpass filtering and motion correction steps, each block of data was truncated, with a pre-stimulus baseline of 3 seconds and a post-stimulus interval of 10 seconds. Additionally, a linear trend was removed from each truncated block signal using the detrend function in MATLAB. Finally, the truncated block time series were averaged across all blocks for each channel, stimulus, and subject, forming the basis for subsequent analysis.

To determine the temporal window for peak hemodynamic activation, we calculated the grand average of all block time series across subjects, channels, and stimuli. We then identified a specific time window following the onset of the stimulus that corresponded to the peak amplitude of the grand average hemodynamic response. On average, the hemodynamic response exhibited a peak response at approximately 8.7 seconds, with a standard deviation of 0.8 seconds.

For the computation of the neurovascular coupling (NVC) parameter, we followed a specific procedure. First, we identified the activation window, which encompassed one second before and after the grand average peak of the hemodynamic response, considering all trials across different stimulus conditions. Within this activation window, we calculated the mean value of the hemodynamic response. Then, we computed the mean value during a pre-stimulus baseline window lasting for 3 seconds. Subsequently, we subtracted the mean value obtained from the pre-stimulus baseline window from the mean value calculated within the activation window.

### 2.6. Data Analysis

All statistical analyses were conducted using JASP software (JASP Team, 2023). To examine the differences between stimulus conditions, a two-way repeated measures ANOVA was conducted for each channel’s group-wise NVC parameter. The analysis included two within-subject factors: valence (positive and negative) and arousal (high and low). Post-hoc t-tests were then performed for each channel data, applying the Bonferroni correction.

To identify the anatomical regions that exhibited significant contrasts, t-values from channels that surpassed the Bonferroni correction threshold were projected onto a standard brain template. This brain template utilized the 10-20 EEG electrode positioning system, enabling the visualization of specific anatomical locations associated with significant differences (Figure 3).

**Figure 3.**
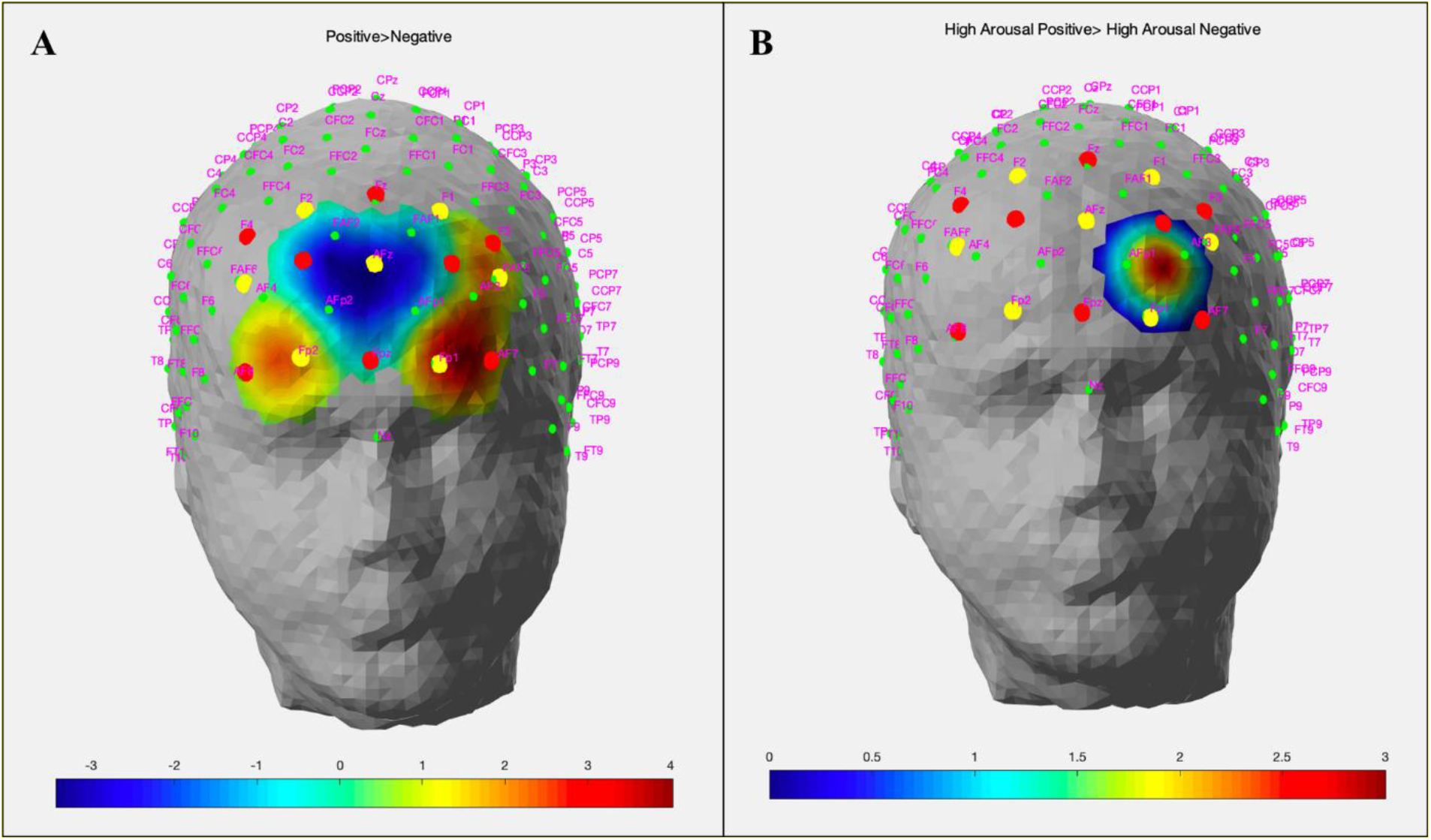
Hemodynamic activity strength parameter map for (A) Positive > Negative, (B) High Arousal Positive > High Arousal Negative conditions. Statistically significant activation channels (p < 0.05) are depicted on a standard head model, along with their thresholded t-statistics scores for each condition.

## 3. RESULTS

The results of the two-way repeated measures ANOVA revealed a significant main effect of valence for channels 10 (F(1,19) = 11.808, p = 0.003), 13 (F(1,19) = 9.842, p = 0.005), 14 (F(1,19) = 5.727, p = 0.027), 17 (F(1,19) = 5.381, p = 0.032), 18 (F(1,19) = 7.884, p = 0.011), and 22 (F(1,19) = 16.192, p < 0.001) while no channels demonstrated a main effect of arousal. An interaction effect between valence and arousal was observed in channel 15 (F(1,19) = 4.837, p = 0.040).

Post hoc tests for channel 15 using Bonferroni correction revealed that high positive stimuli elicited a statistically significant hemodynamic activation than high negative stimuli (p = 0.037). These findings suggest that channel 15, located within BA 10, plays a crucial role in differentiating emotional responses to highly positive and highly negative stimuli (Figure 3B).

Post-hoc tests for channels demonstrating a main effect of valence revealed that regardless of arousal level, all subjects presented a significantly higher hemodynamic activity during processing of positive stimuli in Channels 14 (t=2.393), 17 (t=2.32) and 22 (t=4.024) while they demonstrated a significantly higher hemodynamic activity during processing of negative stimuli in Channels 10 (t=3.436), 13 (t=3.137) and 18 (t=2.808) (Figure 3A).

Regarding the location of channels demonstrating a main effect of valence, we found that channels 10 and 17 were located in the right hemispheric portion of Brodmann area (BA) 10 and 11, respectively. On the other hand, channels 13, 14, and 22 were situated in the left hemispheric portion of BA 10, 46, and 11, respectively.

## 4. DISCUSSION

The primary objective of this study was to assess the viability of utilizing a mobile and wearable fNIRS system to objectively identify the specific regions within the PFC that are involved in the processing of basic emotions. Specifically, we aimed to compare the spatial localization of cerebral activation when individuals were presented with negative and positive valence pictures in the absence of any other interfering stimuli. Our research question centered around whether different basic emotions exhibit recognizable hemodynamic correlates within the PFC that can be measured and identified using fNIRS technology. To address this question, we designed an experimental protocol where healthy participants viewed pictures from the IAPS database, which offers standardized and realistic pictures with pre-determined valence and arousal scores. We selected pleasant and unpleasant pictures with similar valence and arousal scores to examine the hemodynamic activity within the PFC regions during the processing of basic emotions. Our objectives were to investigate the spatiotemporal mapping of PFC activity using fNIRS methodology and to explore the similarities and differences in PFC circuitry activation during the processing of different levels of valence (positive and negative) and arousal (high and low). By employing fNIRS, a wearable, non-invasive, and portable imaging system, we sought to uncover the unique activation patterns associated with different emotional states, thereby advancing our understanding of emotional processing in the PFC.

Prior functional neuroimaging studies have convincingly demonstrated that the human brain consists of intricate neural pathways in the cortical and subcortical regions, which collaborate to handle emotional stimuli. These studies have specifically emphasized the crucial role played by specific subregions within the anterior PFC in tasks related to the processing, assessment, integration, and regulation of emotional information (Damasio A.R., 1996; Davidson et al., 2000). Although fNIRS holds great promise for integration into affective BCI systems, primarily due to its ability to easily acquire functional data from the PFC, it is essential to establish whether the processing of different basic emotions is accompanied by distinctive and independent spatiotemporal patterns of activation within the anterior PFC. Detecting these patterns reliably using fNIRS is a critical initial step in the process of integrating this technology into affective BCIs. This investigation is crucial for validating the effectiveness of fNIRS in capturing emotion-related neural activity and advancing our understanding of the neural mechanisms underlying emotional processing. In the subsequent sections, we delve into a detailed discussion of the neuroanatomical and neurophysiological interpretations of the observed differential activation patterns.

### 4.1. Positive valence stimuli induce higher hemodynamic activation in bilateral orbitofrontal cortex regions when compared to negative valence stimuli

In our study, we observed that positive valence emotional stimuli elicited higher levels of hemodynamic activation in bilateral regions involving the dorsolateral prefrontal cortex (DLPFC) and orbitofrontal cortex (OFC) when compared to the processing of negative emotional stimuli. The DLPFC is known to play a role in regulating both attention and emotion (Golkar et al., 2012). The significant activation of the DLPFC during the processing of negative stimuli may indicate an attentional focus on the presented stimuli. Additionally, the DLPFC is involved in emotion regulation through the reappraisal of the stimuli. Therefore, the bilateral activation of the DLPFC suggests a cognitive demand for regulating positive emotions in response to the stimuli.

The DLPFC and OFC have been recognized as relay centers that integrate emotional, memory-related, and sensory information. This integration of cognitive and emotional processes contributes to the control of emotional reactions and behavioral responses. Neural activity in the OFC is associated with the computation of the motivational and emotional value of presented stimuli (Golkar et al., 2012; Rolls, 2004). The differential activity observed in these regions confirms that our experimental stimuli successfully induced the intended hemodynamic contrast in the expected cerebral regions, validating their use in testing our hypotheses.

### 4.2. Negative valence stimuli induce higher hemodynamic activation in mPFC regions when compared to positive valence stimuli

We found that negative valence emotional stimuli led to higher levels of hemodynamic activation in the medial prefrontal cortex (mPFC) compared to the processing of positive emotional stimuli. The mPFC is implicated in self-referential processing, emotional regulation, and the generation of self-relevant thoughts and emotions (Etkin et al., 2011; Ochsner et al., 2012). The higher activation observed in the mPFC during the processing of negative emotional stimuli may reflect increased engagement of self-referential processing and self-focused rumination (Kross et al., 2009). Negative emotions often prompt individuals to engage in introspection and self-evaluation, potentially leading to the generation of negative self-relevant thoughts. Our findings support the notion that negative valence emotional stimuli may trigger greater involvement of self-related cognitive processes mediated by the mPFC.

### 4.3. High arousal positive valence stimuli induce higher hemodynamic activation in the left DLPFC region when compared to high arousal negative valence stimuli

Our study presents compelling evidence supporting the valence hypothesis, as we observed distinct hemodynamic activity patterns in response to positive and negative valence stimuli while accounting for arousal levels. Specifically, the significantly higher activation of the left DLPFC in the presence of positive valence stimuli, as compared to negative valence stimuli, aligns with Balconi & Mazza’s (2010) proposition that the left PFC is involved in processing positive emotions, while the right PFC is associated with negative emotions. By controlling for arousal levels, we successfully isolated the distinct influence of valence on brain activity, further supporting the notion of lateralized emotional processing within the prefrontal cortex. These findings significantly contribute to our understanding of the neural mechanism of emotional experiences, highlighting the pivotal role of the left DLPFC in mediating positive emotional responses and advancing our broader comprehension of emotion-related neural mechanisms.

## 5. CONCLUSION

The study presented here employed fNIRS technology to investigate the engagement of PFC regions in the processing of basic emotions. Through the analysis of spatiotemporal patterns of hemodynamic activity, the research aimed to discern specific cortical responses associated with different emotional states. The results revealed distinct hemodynamic activity patterns in the PFC region corresponding to different basic emotions, thereby supporting the feasibility of utilizing mobile fNIRS for objective and real-time decoding of emotional states. This advancement holds promise for various applications in emotion recognition and related fields.

Furthermore, the study’s outcomes suggest that integrating fNIRS-derived biological features with multivariate pattern analysis and machine learning algorithms could enhance the real-time detection of emotional states. By combining these approaches, it becomes possible to identify the neuronal activation patterns underlying various emotional states more accurately in real-time scenarios.

Overall, the observed distinct hemodynamic patterns in response to different emotional stimuli showcase the potential of fNIRS technology for objectively identifying and characterizing neuronal activation patterns associated with diverse emotional states. This progress opens up new avenues for understanding the neural basis of emotions and contributes to the development of innovative tools for emotion-related research and applications.

